# RAS specific protease Induces Irreversible Growth Arrest via p27 in several KRAS Mutant Colorectal Cancer cell lines

**DOI:** 10.1101/2020.12.08.416578

**Authors:** Caleb K. Stubbs, Marco Biancucci, Vania Vidimar, Karla J. F. Satchell

**Affiliations:** Department of Microbiology and Immunology, Northwestern University, Feinberg School of Medicine, Chicago, Illinois

## Abstract

Ras-specific proteases to degrade RAS within cancer cells are under active development as an innovative strategy to treat tumorigenesis. The naturally occurring biological toxin effector called RAS/RAP1-specific endopeptidase (RRSP) is known to cleave all RAS within a cell, including HRAS, KRAS, NRAS and mutant KRAS G13D. In the course of studies developing RRSP as an anti-cancer therapeutic, it was shown that cleavage of total RAS by RRSP results in a range of cell fates from cytotoxicity to moderate growth inhibition. Despite the considerable amount of evidence demonstrating RRSP anti-tumor effects *in vivo,* our understanding of the mechanisms involved are unknown. Here, we first demonstrate, using isogenic mouse fibroblasts expressing a single isoform of RAS or mutant KRAS, that RRSP equally inactivates all isoforms of RAS as well as the major oncogenic KRAS mutants. The cleavage of RAS inhibited phosphorylation of ERK and cell proliferation regardless of the RAS isoform. To investigate further how RAS processing might lead to varying outcomes in cell fate within cancer cells, we tested RRSP against four colorectal cancer cell lines with a range of cell fates. While cell lines highly susceptible to RRSP (HCT116 and SW1463) undergo cytotoxic death, RRSP treatment of GP5d cells induces G1 cell cycle arrest, and SW620 cells instead induces growth inhibition through cell senescence. In three of four cell lines tested, growth effects were dictated by rescued expression of the tumor suppressor protein p27 (Kip1). The ability of RRSP to inactivate all RAS and inhibit cancer cell growth through a variety of mechanisms highlights the antitumor potential of RRSP, and further warrants investigation as a potential anti-tumor therapeutic.

## Introduction

The oncoprotein Rat sarcoma (RAS) GTPase cycles between GTP-bound (active) and GDP-bound (inactive) states for activation of downstream effectors, each playing key roles in cell proliferation and survival (1). This process is highly reliant on GTPase activating proteins (GAPs) and guanine exchange factors (GEFs) for hydrolysis of GTP and nucleotide exchange of GDP to GTP, respectively (2, 3). Upon growth receptor stimulation, activated RAS recruits downstream effectors, including Rapidly Accelerated Fibrosarcoma kinase and phosphatidylinositol-3-kinase. These effectors subsequently activate signaling pathways responsible for cell growth and survival, including the mitogen-activated kinase (MEK) to extracellular signal-regulated kinase (ERK) signaling pathway and the protein kinase B (also known as AKT) to mammalian target of rapamycin (mTOR) pathway, respectively (4, 5).

Thirty percent of all human cancers contain mutations in *RAS* (1,6). Mutant *RAS,* paired with loss of function in tumor suppressor genes such as *TP53* and *APC,* are sufficient to fully transform cells and drive tumorigenesis (6). Nearly all RAS mutations occur as point mutations at Gly-12, Gly-13 or Gln-61, resulting in constitutive activation of RAS (1). Among the major *RAS* isoforms *(HRAS, NRAS,* and *KRAS), KRAS* is the most frequently mutated isoform among all cancers (85%) followed by *NRAS* (11%) and *HRAS* (4%) (6). *RAS* mutations are highly enriched specifically in three of the four most lethal cancers in the United States, including pancreatic adenocarcinoma (98%), colorectal adenocarcinoma (52%), and lung adenocarcinoma (32%) (1, 6).

Although numerous studies support the advantages of targeting RAS to treat cancer, it remains an unsolved challenge in the clinic (7–11). Recent studies have taken advantage of biochemical properties of specific RAS mutants to develop selective small molecule inhibitors specific for highly oncogenic mutant forms of RAS. In particular, small molecules targeting KRAS G12C have been developed and are undergoing clinical trials (12–14). Despite this success, the strategy of selective inhibition has problems of being applicable to only a limited range of cancers integrated with personalized medicine and cannot be used to treat cancers that lack the specific mutation. To address this gap, new approaches are being developed to more broadly target RAS either with proteases that directly cleave RAS (15, 16) or with linkers that target RAS for cellular degradation (17–20).

Our lab has identified a potent protease that cleaves RAS called the Ras/Rap1-specific endopeptidase (RRSP). RRSP is a small domain of a large toxin secreted by the bacterium *Vibrio vulnificus* during host infection. *V. vulnificus* delivers RRSP into intestinal epithelial cells during host infection, where it targets all RAS isoforms and close homolog Ras-related protein 1 (RAP1). Through RAS inactivation, this bacterium suppresses the host immune response, thereby aiding systemic dissemination of the bacterium (21, 22). Detailed structural and biochemical studies have shown that RRSP attacks the peptide bond between Tyr-32 and Asp-33 in the Switch I region of both RAS and RAP1 (23). As a result, RAS and RAP1 are unable to undergo GTP-GDP exchange or bind to their downstream effectors (24, 25). Recently, RRSP engineered as a chimeric toxin for *in vivo* delivery was shown to significantly reduce breast and colon tumor growth in xenograft mouse models (15).

The potential applicability of RRSP to a broader range of cancers was tested *in vitro* using the standardized National Cancer Institute (NCI) cancer cell panel (26). Fourteen of 60 cell lines were classified as highly susceptible with cells undergoing cytotoxic cell death. Further, 38/60 of cell lines showed significant growth inhibition. Only 8/60 showed low or no susceptibility with cell growing near normal rates, possibly due to lack of the receptor for the engineered chimeric toxin (15). The range of cell fates highlights that the cellular responses to total RAS cleavage are variable and the mechanism by which RRSP inhibits growth inhibition is currently unknown. Here, we investigate how RRSP processing affects cell signaling and characterize the impact of cleaving total RAS on cancer cell growth and survival. We further demonstrate that RRSP can disrupt colorectal cancer (CRC) cell growth through multiple mechanisms, including loss of cell viability, cell cycle arrest, and senescence.

## Results

### RRSP cleaves and inhibits proliferation in RAS wildtype and KRAS mutant cells

RRSP was previously shown to specifically cleave HRAS, NRAS, and KRAS when the proteins were ectopically expressed in HeLa cells and recombinant RRSP was shown to process purified KRAS G12D, G13D, and Q61R in biochemical assays (23). To get a broader sense of RRSP effectiveness across different isoforms and mutants of RAS, we tested RRSP against the ‘RAS-less’ mouse embryonic fibroblast (MEF) cell line panel developed by Drosten et al. (5). These isogenic cell lines have endogenous *RAS* genetically deleted from their genome and replaced with a single allelic copy of human *RAS* gene. For delivery of RRSP into mouse cells, we used the anthrax toxin-based delivery system wherein the anthrax toxin lethal factor N-terminus was fused with RRSP (LF_N_RRSP) or LF_N_RRSP with a catalytically inactivating H4030A amino acid substitution (here after referred to as LF_N_RRSP*). Intracellular delivery of RRSP (previously known as DUF5) with this system has been previously been demonstrated in several mammalian and mouse cell lines (23, 27, 28).

In MEFs expressing human KRAS, HRAS, or NRAS, treatment with 3 nM LF_N_RRSP dramatically decreased intact full-length RAS levels with increased detection of cleaved RAS. For each isoform, LF_N_RRSP was found to cleave at least 80% of RAS after 24 hours (**Fig. 1A)**. As expected, controls treated with PA alone or in combination with catalytically inactive LF_N_RRSP* showed no change of intact RAS protein levels **(Fig. 1A and 1B).** We also observed similarities of RRSP activity in MEF cell lines expressing oncogenic KRAS, including G12V, G12D, G12C, G13D, and Q61R. Amongst each of the mutant RAS alleles tested, we observed ~25% of total RAS remaining following LF_N_RRSP treatment, with no significant loss of RAS in cells treated with PA (**Fig. 1B).** The oncogenic RAS variants with the higher percentage of RAS remaining following LF_N_RRSP treatment were G13D, G12C and Q61R, although these differences were not statistically significant. Further, the total RAS remaining in each LFNRRSP-treated MEF cell line was not statistically significant between groups. In addition to cleavage of RAS, RRSP treated cells showed significant decreases in phosphorylation of ERK when compared to cells treated with PA alone or with the catalytically inactive LF_N_RRSP* (**Fig. 1C and D)**.

**Figure 1.**
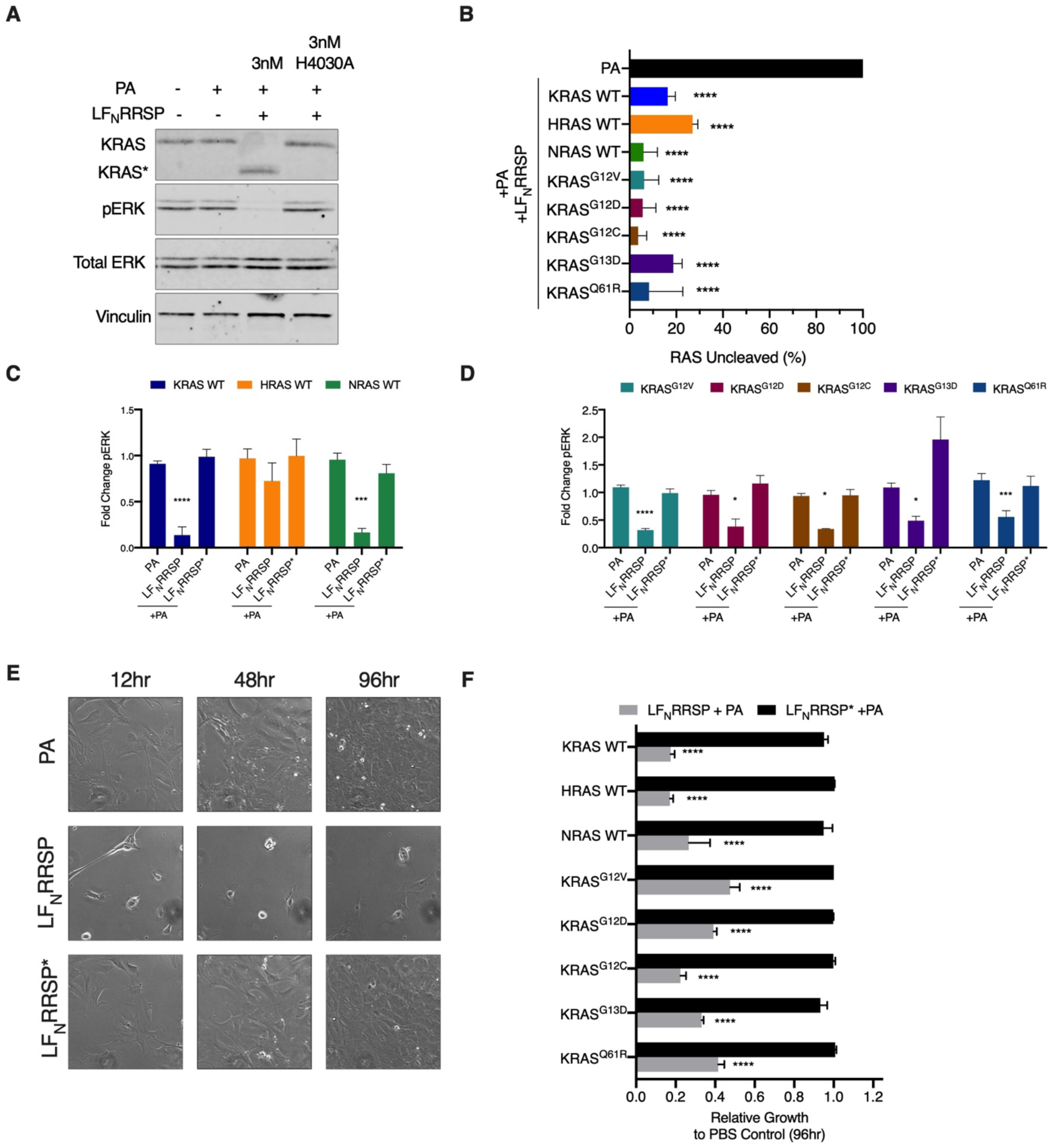
LF_N_RRSP cleaves and inhibits all RAS isoforms and KRAS oncogenic mutants in RAS- less MEFs. (**A**) Representative western blot analysis of LF_N_RRSP cleavage of RAS and inhibition of ERK in KRAS WT RAS-less MEFs after 24 hr. Vinculin was used as a gel loading control. (**B**) Densitometric analysis of total percent RAS following LF_N_RRSP treatment after 24 hr in RAS-less MEF cell lines; *n* = 3. (**C and D**) Densitometric analysis of fold change in pERK compared to PBS control after 24 hr for RAS-less MEF cell lines described; *n* = 3. (**E**) Brightfield images of KRAS WT RAS-less MEFs treated with either PA alone or in combination with LF_N_RRSP or LF_N_RRSP* at indicated timepoints. (**F**) Relative growth inhibition in RAS-less MEFs compared to PA control at 96 hours following treatment with either LF_N_RRSP or LF_N_RRSP*. Results are expressed as mean ± SEM of three independent experiments (*P<0.05, **P<0.01, ****P<0.0001 versus PA control as determined through one-way ANOVA followed by Dunnett’s multiple comparison test).

To test the impact of processing of different RAS isoforms on cell proliferation, RRSP- treated cells were tracked using time lapse imaging for four days. At early timepoints following treatment, LFNRRSP-induced a severe cell rounding that was not observed in PA alone and LF_N_RRSP* control treated cells (**Fig. 1E)**. This phenotype is consistent with previous studies with RRSP and is possibly linked to cleavage of RAP1, which regulates cytoskeletal dynamics (23, 27, 29). Across all MEF cell lines, LF_N_RRSP inhibited growth by at least 60% compared to PA only and LF_N_RRSP* mutant controls (**Fig. 1F, Supplementary Fig. 1**).

Altogether, these results in MEFs demonstrate that RRSP is equally able to cleave all isoforms of RAS and mutant KRAS to inhibit both ERK phosphorylation and cell proliferation in a defined system. Thus, differences in cancer cell fate upon treatment with RRSP may depend on processes downstream of RAS processing that could vary in different cell lines.

### RRSP inhibits proliferation and pERK activation in CRC cell lines

To probe further the effect of processing of RAS on downstream signaling, we focused on four KRAS mutant CRC cell lines (**Fig. 2A**). Due to problems with variable expression of the anthrax toxin receptor on the selected human cancer cells, we switched to a recently described, highly potent RRSP chimeric toxin wherein RRSP is tethered to the translocation B fragment of diphtheria toxin (RRSP-DT_B_) (15). Similar to the anthrax toxin system, RRSP-DT_B_ binds to a human receptor (heparin binding epidermal growth factor-like growth factor (HB-EGF)), is endocytosed, and translocated into the cytosol across the vacuolar membrane. Expression of HB- EGF receptor was found to be similar between the selected CRC cell lines (**Supplementary Fig. 2A**).

**Figure 2.**
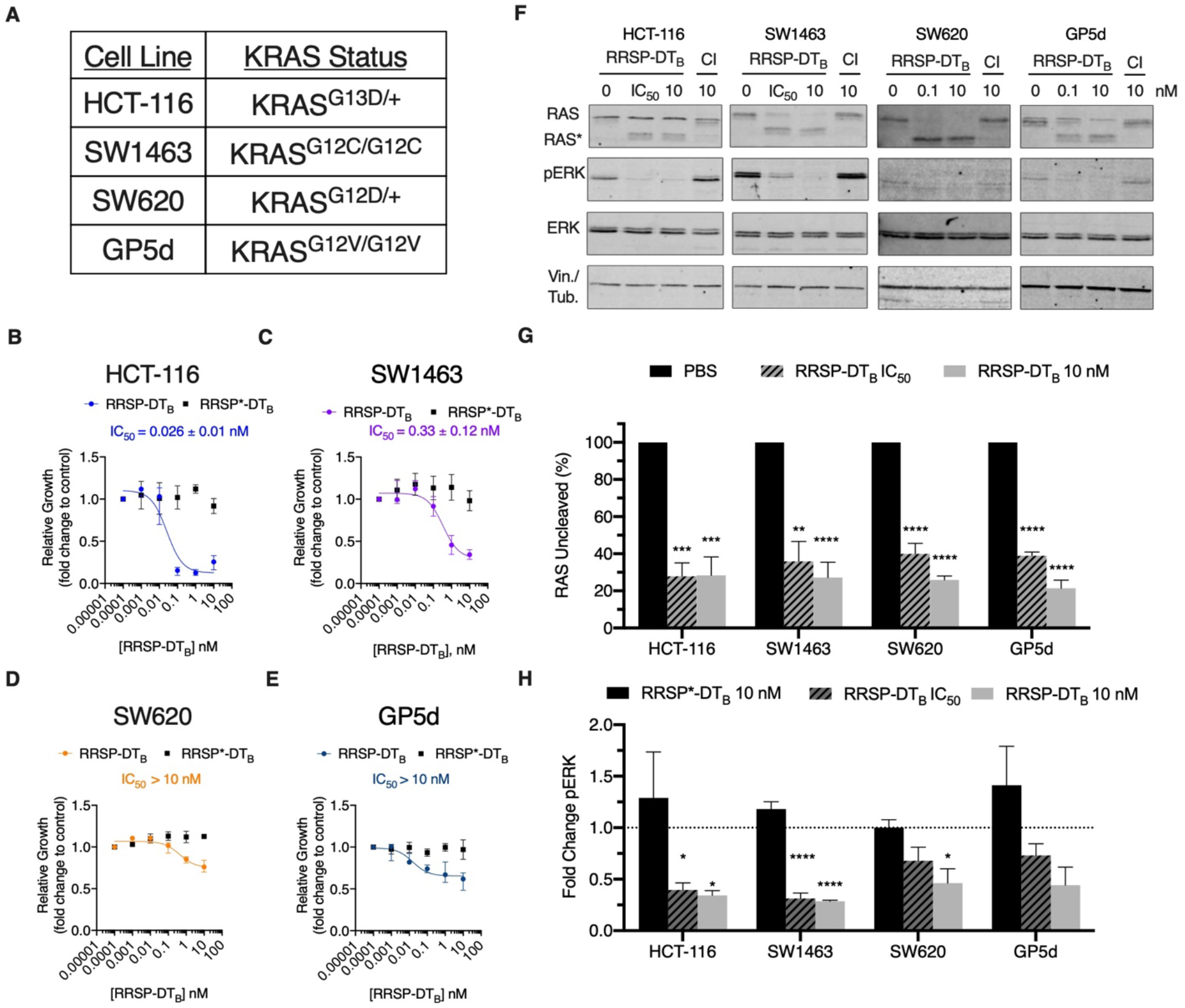
RRSP-DT_B_ growth inhibition in CRC cell lines as defined by (36). (**A**) Cell line panel of KRAS mutant CRC cells. (**B-E**) Fitted dose response curve of RRSP-DT_B_ in CRC cell lines after 24 hr. Results are displayed as mean ± SEM, *n* = 4. (**F**) Representative western blot analysis of RAS cleavage and ERK inhibition in CRC cell lines treated with either RRSP-DT_B_ or catalytically inactive mutant (labeled by CI) after 24 hr. All concentrations are expressed in nanomolar. In all cell lines, vinculin was used as gel loading control except SW620 cells in which aTubulin was used. (**G** and **H**) Densitometric analysis of fold change in percent total RAS and pERK compared to PBS control after 24 hours in CRC cell lines; *n* = 3. Results are expressed as means ± SD of three independent experiments (*P<0.05, **P<0.01, ****P<0.0001 versus PBS control as determined through one-way ANOVA followed by Dunnett’s multiple comparison test).

To examine RRSP growth sensitivities between the CRC cell lines, cells were treated with increasing concentration of RRSP-DT_B_ or with catalytically inactive RRSP-DT_B_ (RRSP*-DT_B_) and growth inhibition was monitored. HCT-116 cells showed the greatest sensitivity to loss of RAS due to RRSP in time lapse video microscopy (**Fig. 2B, Supplementary Fig. 2B and C**), consistent with prior data that HCT-116 are highly susceptible to RRSP (15). Cells treated with catalytically inactive RRSP*-DT_B_ showed no difference, confirming the sensitivity was due to processing of RAS (**Fig. 2B, Supplementary Fig. 2B-E and L**). A similar effect was observed in SW1463 cells, where RRSP-DT_B_ severely inhibited growth at each time point (**Fig. 2C, Supplementary Fig. 2D-E**), revealing this CRC cell line is also highly susceptible. By contrast, cell lines SW620 and GP5d were less susceptible, and yet showed growth inhibition when compared to cells treated with the catalytically inactive RRSP*-DT_B_ (**Fig. 2D and 2E, Supplementary Fig. 2F-I**). Across all of the cell lines, at least 80% of total RAS was cleaved by RRSP (**Fig. 2F and 2G**). In addition, phosphorylation of ERK was significantly reduced compared to respective RRSP*-DT_B_ treated samples (**Fig. 2F and 2H**).

These differences in growth following RRSP treatment further impacted long term survival. In highly susceptible cell lines HCT-116 and SW1463 cells, RRSP was cytotoxic with a much lower relative percent cell viability compared to mock treated controls (**Fig. 3A**). By contrast, GP5d and SW620 were not affected in cell viability. Yet, when SW1463 and GP5d cells were treated for 48 hours and then reseeded at low cell densities, both showed a decrease in colony formation, suggesting that RRSP can induce a permanent non-proliferative state (**Fig. 3B and 3C**). SW620 cells also showed significant increase activity of the enzyme ß-galactosidase, a marker of senescence marker. However, ß-galactosidase activity of treated GP5d cells remained unchanged (**Fig. 3D**). Altogether, these data demonstrate that RRSP is cytotoxic in some highly susceptible cell lines, while in less sensitive cell lines the cells remain metabolically active, but are unable to proliferate and, in some cases, enter into senescence.

**Figure 3.**
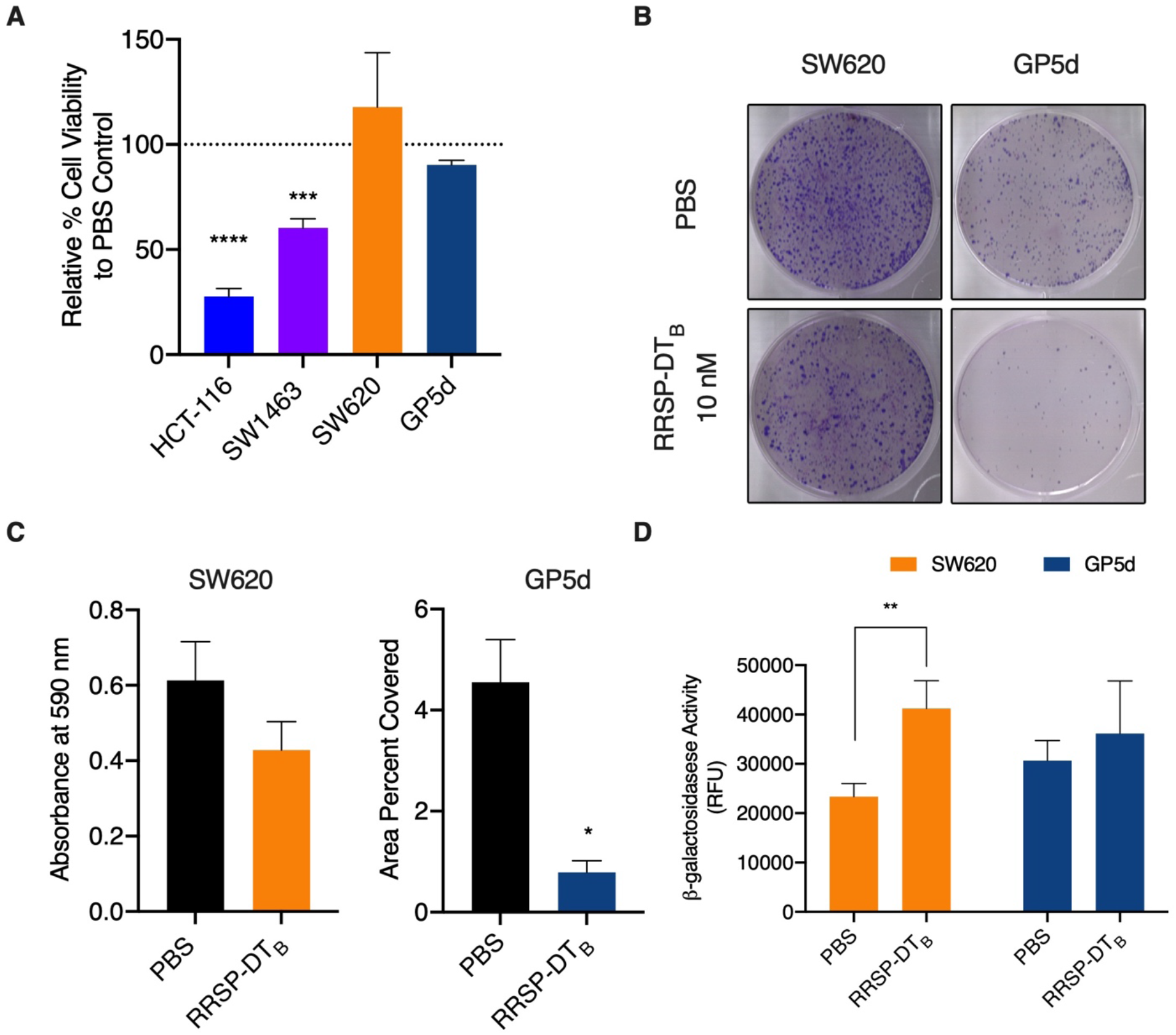
RRSP-DT_B_ decreases viability and causes irreversible growth inhibition in KRAS dependent and independent CRC cell lines. (**A**) Relative percent viability after 72-hour treatment with 10 nM RRSP-DT_B_ compared to PBS control in CRC cell lines. (**B**) Representative images of crystal violet-stained colonies from RRSP less sensitive cell lines pretreated with 10 nM RRSP-DT_B_ for 48 hours and replated at low seeding density to form colonies over 14 days. (**C**) Quantitative analysis of crystal-violet stained colonies from less sensitive RRSP cell lines from three independent experiments. Results are expressed as means ± SD of three independent experiments (**D**) Measured cell senescence activity in RRSP less sensitive cell lines treated with 10 nM RRSP-DT_B_ for 48 hours then incubated with SA-ß-Gal Substrate for 1 hour at 37°C, *n* = 3. All results described above are expressed as mean ± SEM of three independent experiments (*P<0.05, **P<0.01, ****P<0.0001 versus PBS control as determined through one-way ANOVA followed by Dunnett’s multiple comparison test).

### RAS cleavage can induce upregulation of cyclin-dependent kinase inhibitor p27 and hypophosphorylation of RB

We next took advantage of the unique cell line specific effects on cell growth and survival to better understand the underlying mechanisms regulating cell fate following RAS inhibition. Cell lysates from treated or untreated HCT-116 (highly sensitive) and SW620 (less sensitive) were incubated overnight with nitrocellulose membranes containing capture antibodies towards 43 different phosphorylated proteins (**Fig. 4A**). For RRSP-treated HCT-116 cells, there was increased phosphorylation observed for cell stress proteins such as p38α, p90 ribosomal S6 kinase (RSK1/2/3), and Jun-activated kinase (JNK) (**Fig. 4B, Supplementary Fig. 3**). In addition, RRSP treatment increased phosphorylation of several Signal Transducer and Activator of Transcription (STAT) transcription factors. By contrast, the less responsive SW620 cells showed decreased phosphorylation of several STAT proteins (**Fig. 4B, Supplementary Fig. 3**). We also observed a significant fold increase in With No K(lysine)-1 (WNK1) kinase at Thr-60. This kinase is phosphorylated by AKT in HEK293 cells and is best known for regulating ion transport across membranes (30). However, phosphorylation of Thr-60 has no effect on its kinase activity or its cellular localization (31). Because RRSP decreases AKT activation (**Fig. 4B**), it is unlikely that WNK1 Thr-60 phosphorylation is involved in the growth differences we observe between cell lines.

**Figure 4.**
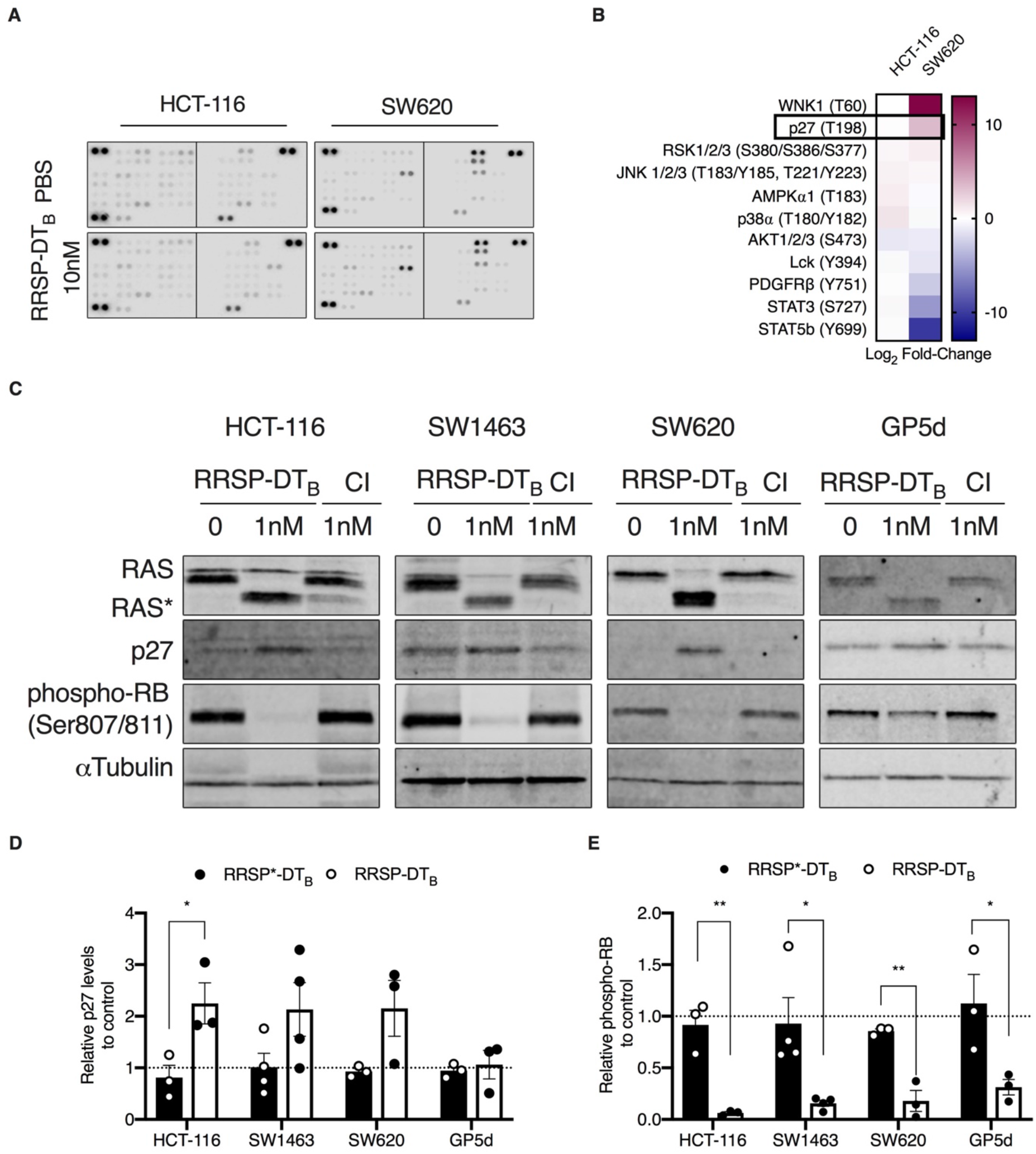
RRSP-DT_B_ cleavage of RAS induces p27 expression in CRC cell lines. (**A**) Human phospho-kinase array blots of HCT-116 and SW620 cells treated with either PBS or RRSP-DT_B_ (10nM) for 24 hours. (**B**) Densitometric analysis kinase array depicted through a heatmap of relative phosphorylated proteins levels in response to RRSP-DT_B_ compared to PBS control in HCT-116 and SW620 cells, *n* = 1. (**C**) Representative western blot images of p27 and phosphor- RB expression in CRC cell lines treated with either RRSP-DT_B_ or RRSP*-DT_B_ for 24 hours. (**D** and **E**) Densitometric analysis of fold change in p27 and phospho-RB compared to PBS after 24 hr in RRSP-DT_B_ treated CRC cell lines; *n* = 3. aTubulin was used as gel loading control. Results are expressed as mean ± SEM of three independent experiments (*P < 0.05, ** < 0.01, **** < 0.0001 versus PBS control as determined through one-way ANOVA followed by Dunnett’s multiple comparison test).

Thus, we focused on the large fold-change difference observed in phosphorylation at Thr- 198 of the cyclin-dependent kinase inhibitor p27 (also known as Kip1) (**Fig. 4B**). RAS is known to regulate critical components involved in cell cycle. RAS activation is directly linked to hyperphosphorylation of retinoblastoma protein (RB), thereby relieving its repression of E2F transcription factors, allowing transcription of G1 promoting genes, and promoting the cell cycle to progress from G1 to S phase (32). Previous studies have established that phosphorylation of p27 at Thr-198 is critical for stabilizing p27 expression by preventing ubiquitin-dependent degradation (33). In fact, aberrant RAS activity in cancer cells causes p27 post-translational downregulation through both ERK and AKT (34–36). These data support that inhibition of RAS by RRSP could lead to downstream rescue expression of p27 expression in cells, thereby slowing reversing the hyper-phosphorylation of RB.

To test this possibility, all four cancer cell lines were treated with a sublethal dose of RRSP-DT_B_. The treatment increased p27 protein levels in HCT-116, SW620, and SW1463 cells, while in GP5d cells levels remained unchanged (**Fig. 4C-D**). Concomitant with upregulation of p27, all cell lines showed a significant decrease in RB phosphorylation at Ser-807/Ser-811 (**Fig. 4C and 4E**). Unfortunately, total RB was undetectable using commercially available antibodies. To be confident that RB hypo-phosphorylation was not due to low RB expression, we transiently expressed green-fluorescent protein (GFP)-tagged RB in HCT-116 cells (**Supplementary Fig. 4A**). In GFP-RB expressing cells treated with RRSP-DT_B_, hypo-phosphorylation of RB protein compared to PBS and RRSP*-DT_B_ controls was observed (**Supplementary Fig. 4B**). Protein levels of GFP-RB decreased in RRSP-DT_B_, consistent with a role of p27 in regulating RB expression (37).

Unexpectedly, hypo-phosphorylation of RB was also observed in GP5d cells despite showing no change in the expression or phosphorylation of p27. The cyclin-dependent kinase inhibitor, p21, also plays a critical role in RB regulation. However, there was also no change in p21 protein levels in RRSP-treated GP5d cells (**Supplementary Fig. 4C**).

### RRSP induces G1 phase cycle arrest

Elevated p27 protein expression in combination with hypo-phosphorylation of RB suggested that RRSP treatment induces cell cycle arrest. Under normal conditions, p27 regulates G1 checkpoint during the cell cycle by preventing entry into S phase through inhibition of CDK2 (38, 39). To test if RRSP-DT_B_ treatment induces cell cycle arrest, cell lines were treated for 24 hours and the percentage of cells in G1, S, or G2/M phase was monitored. all cell lines that showed reduced RB phosphorylation had significant population of cells locked in the G1 state compared to PBS and RRSP*-DT_B_ treated samples (**Fig. 5, Supplementary Fig. 6**). The most dramatic increase in G1 arrest was seen in SW620 cells, where nearly 100% of cells remained in the G0/G1 phase following RRSP-DT_B_ treatment (**Fig. 5C**). This G1 cell arrest was dependent of the RAS processing activity of RRSP as the catalytically inactive mutant RRSP-DT_B_ did not induce the cell cycle arrest (**Fig. 5A-D**). Together, these data illustrate that RRSP cleavage of RAS can induce growth inhibition through several different mechanisms, some of which are still only partially understood.

**Figure 5.**
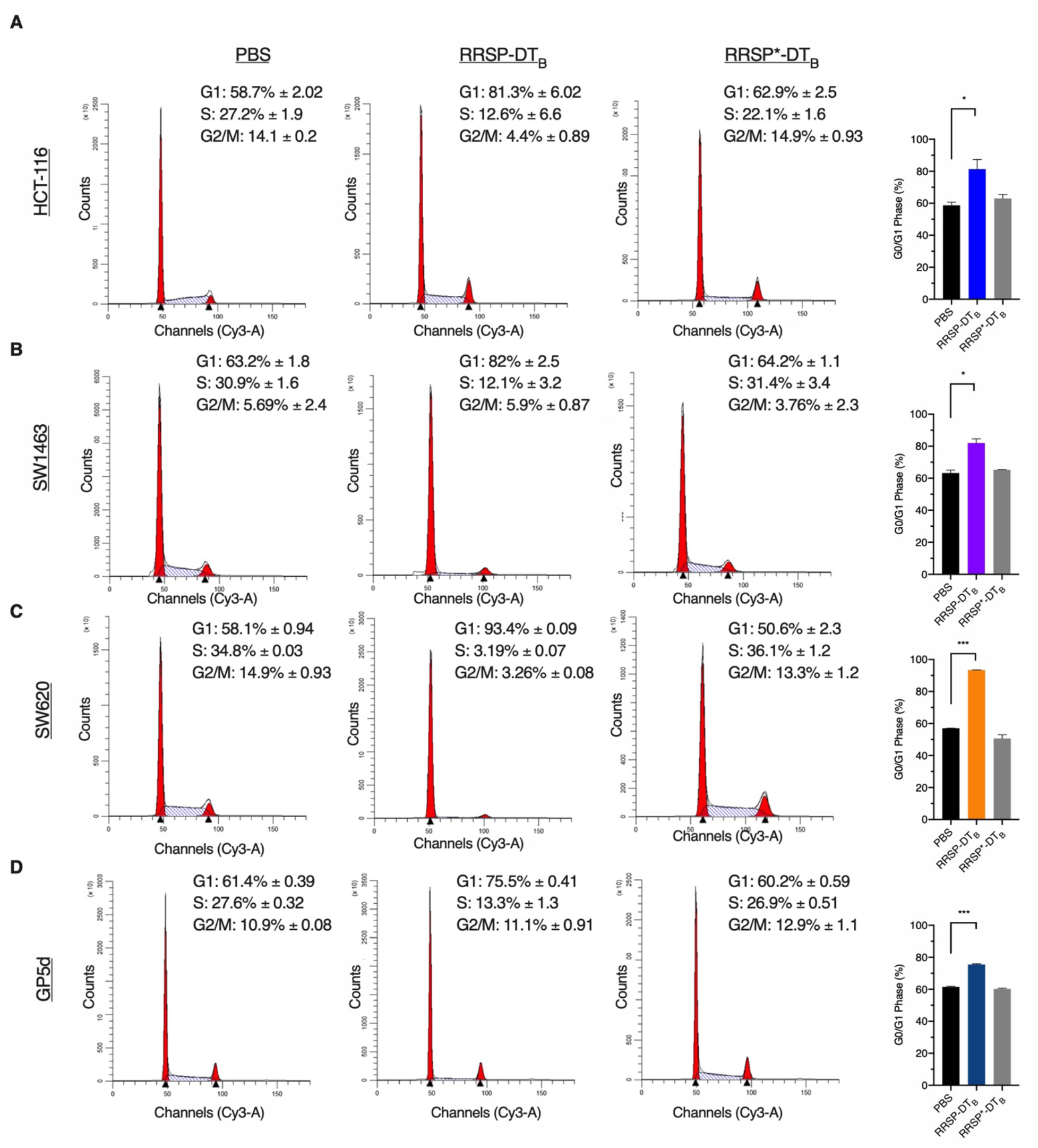
RRSP-DT_B_ induce G1 cell cycle arrest in CRC cell lines. (**A-D**) Cell cycle flow cytometry analysis of CRC cell lines treated with either PBS, RRSP-DT_B_ or RRSP*-DT_B_ (1 nM) for 24 hours. Percentage of cells in each phase are an average of three independent experiments. Bar graphs depict percentage of cells in G1 phase for each treated sample; *n* = 3. Results are expressed as mean ± SEM of three independent experiments (*P<0.05, **P<0.01, ****P<0.0001 versus PA control as determined through one-way ANOVA followed by Dunnett’s multiple comparison test).

## Discussion

It has been over 30 years since the discovery of the importance of RAS for driving tumorigenesis in cancer. Lung, pancreatic, and colorectal cancers remain being the most lethal cancers in the United States with high mutation rates in RAS, the most commonly mutated isoform of KRAS. Despite the significant amount of research being conducted on RAS, it still remains a challenging target in the field. Small molecules directed to specific RAS mutants, specially KRAS G12C, have shown promising results in clinical trials (40), but will only benefit a small subset of patients. Our lab has discovered RRSP as a potent, site specific inhibitor of RAS capable of inhibiting all RAS isoforms simultaneously along with downstream activation. RRSP antitumorigenic effects are well demonstrated *in vivo* with xenograft models for both breast and colon cancers, wherein tumor growth was stunted and, in some cases, showed regression (15). Evidence for RRSP as a therapeutic inhibitor of RAS is sufficient, however the mechanism by which RRSP mediates growth inhibition has been an outstanding question. In this study, we examined the signaling consequences of cleavage of all RAS in several CRC cell lines and its downstream implications on cell proliferation and survival.

First, we examined whether RRSP was a suitable inhibitor across RAS variants. Using the isogenic ‘RAS-less’ MEF model, we demonstrated that all three major RAS isoforms and frequently observed KRAS mutants were equivalent substrates for RRSP. Loss of RAS resulted in reduced ERK activation, which as expected, negatively affected proliferation. Most importantly, we observed no significant differences in RAS cleavage between wildtype isoforms and KRAS mutants.

We next examined RRSP effectiveness in four CRC cell lines, which displayed variations in susceptibility to RRSP. Two of the cell lines with the greatest RRSP growth sensitivity, HCT- 116 and SW1463, had dramatically lower metabolic activity compared to controls indicting cytotoxicity in response to RRSP. Interestingly, GP5d and SW620 retained normal metabolic activity, yet showed significant inability to form colonies following RRSP treatment, mimicking a senescent-like phenotype. From the two cell lines tested, only SW620 had elevated ß- galactosidase activity indicting RRSP could also activate senescent-like mechanisms.

Mechanisms that link RAS and the cell cycle have been well examined. In quiescent cells, p27 is highly expressed in order to inhibit CDKs activity and to suppress RB phosphorylation (38, 39). Upon mitogen stimulation, RAS activation suppresses p27 protein expression through post-translational modifications that signal for its ubiquitin-mediated degradation (34–36). In RAS-driven human cancers, low levels of p27 are frequently observed.

We demonstrated that the growth inhibition in HCT-116, SW1463, and SW620 is the result of G1 cell cycle arrest through the upregulation of p27. These data suggest that RAS cleavage in certain CRC cell lines induces p27 upregulation, leading to a cell cycle arrest state that can induce cell death at prolonged timepoints. Transient overexpression of p27 is known to then induce cell cycle arrest and later apoptosis (41, 42). Although in our studies, only the highly susceptible cells undergo cell death, whereas SW620 retains metabolic activity and undergoes senescence.

Unexpectedly, GP5d cells did not show upregulation of p27 or p21, although RB was hypo- phosphorylated and cells initiated G1 cell cycle arrest. These data support there is another path to growth inhibition induced by RRSP that is not yet understood. In fact, RRSP from the insect pathogen *Photorhabdus asymbiotica,* which is %identical to RRSP, also cleaves RAS (23) and was recently reported to induce G1 cell cycle arrest (43). The proposed mechanism did not depend on RAS processing, and instead involved RRSP directly binding to CDK1 when it is overexpressed, essentially functioning as a protein trap. Thus, the multi-domain RRSP may possess at least two mechanisms to inhibit the cell cycle, one by restoring p27 downstream of RAS processing and another by directly binding to CDK1. Notably, because low p27 expression levels have been correlated with poor survival in patients with different types of cancer including colon, the ability of RRSP to restore p27 expression and to initiate cell cycle arrest could have important implications for the treatment of tumors with aberrant RAS signaling.

## Materials and Methods

### Cell Lines

‘RAS-less’ mouse embryonic fibroblast (MEF) cells were provided by the NCI RAS Initiative at Frederick National Laboratory for Cancer Research (FNLCR). HCT-116 cells were purchased from the American Type Culture Collection. SW1463, GP5d, and SW620 were provided by the NCI. Each cell line was validated by the Northwestern University Sequencing Core by Short Tandem Repeat profiling.

All cells were cultured at 37°C and 5% CO_2_ atmosphere. HCT-116, SW1463, GP5d, SW620 cells were cultured in Dulbecco’s Modified Eagle Medium (DMEM)-F12 with Glutamax (Gibco) containing 10% fetal bovine serum (FBS; Gemini Bio) and 1% penicillin/streptomycin (P/S; Invitrogen). All MEF cells, except for HRAS RAS-less MEFs, were cultured in DMEM (Gibco) with 10% FBS,1% P/S, and 4 μg/ml of blasticidin (ThermoFisher Scientific). HRAS RAS- less MEFs was cultured in 2.5 μg/ml of puromycin (ThermoFisher Scientific).

### Antibodies

Anti-RAS monoclonal antibody recognizing G-domain of all major RAS isoforms was purified from a hybridoma cell line provided by FNLCR and used at 1:2000 dilution as previously described (15). Other commercially available primary antibodies used were: anti-Phospho-p44/42 MAPK (phosphorylated ERK1/2, Thr202/Tyr204, Cell Signaling Technology #4377), anti-p44/42 MAPK (ERK1/2, Cell Signaling Technology #4696), anti-HB-EGF (R&D Systems, #AF-259-NA;), anti- p27^Kip1^ XP (Cell Signaling Technology #3686), Phospho-RB (Ser807/Ser811, Cell Signaling Technology #8516), anti-p21^WAF/Cip1^ (Cell Signaling Technology #2947T), anti-α-Tubulin (Cell Signaling Technology #2144), and anti-vinculin (Cell Signaling Technology #13901). Fluorescently-labeled secondary antibodies obtained from LI-COR Biosciences and used at 1:10,000 dilution were: IRDye 680RD goat anti-mouse (926–68070), IRDye 800CW goat antirabbit (925–322211), and IRDye 800CW donkey anti-goat (925–32214). Western blot images were acquired using an Odyssey Infrared Imaging System (LI-COR Biosciences) and quantified by densitometry using NIH ImageJ software.

### pRB-GFP Transfection

Plasmid RB-GFP FL for expression of GFP-tagged RB was obtained from Addgene (Catalog #16004). For ectopic gene expression, cell lines were transfected using FuGene HD (Promega) according to the manufacturer’s protocols. GFP fluorescence was analyzed using EVOS XL Core imaging system.

### Western Blotting

Cells were washed in PBS and then resuspended in lysis buffer [150 mM NaCl, 20 mM Tris (pH 7.5), 1% Triton X-100, and Pierce Protease phosphatase inhibitor (Sigma-Aldrich)]. Lysates were incubated for 15 minutes on ice and centrifuged at 20,000 x *g* at 4°C for 15 minutes. The concentration of protein in the collected supernatant fluid was determined using the bicinchoninic acid (BCA) assay (ThermoFisher Scientific, no. 23227). Samples were boiled at 95°C in Laemmli SDS loading buffer for 10 minutes and protein was separated on either 15 or 18% SDS-polyacrylamide gels. Proteins were transferred to nitrocellulose membranes and blocked in Trisbuffered saline (TBS) [10 mM Tris (pH 7.4) and 150 mM NaCl] with 5% (w/v) milk for 1 hour.

Membranes were washed with TBS and then incubated in indicated primary antibodies in TBS with 5% (w/v) Fraction V bovine serum album (Fisher BioReagents #194850) overnight at 4°C. Total percentage RAS was calculated using the following equation: % Total RAS = uncleaved RAS band / (RAS uncleaved band / RAS cleaved band) X 100.

### Purifiacation and Intoxication of LF_N_RRSP in MEFs

Recombinant LF_N_RRSP and LF_N_RRSP* were expressed in *Escherichia coli* BL21(DE3) and purified over a HisTrap FF nickel affinity column followed by Superdex 75 size exclusion chromatography using the ÄKTA protein purifier purification system (GE Healthcare), as previously described (27). For intoxication, MEFs were seeded in 6-well plates at 3 x 10^5^ cells per well for 1 hour, after which medium was replaced with fresh medium containing with 7 nM Protective antigen (PA) alone (List Labs, #171E) or in the presence of 3 nM LF_N_RRSP/ LF_N_RRSP^H4030A^ and incubated at indicated timepoints at 37°C in the presence of 5% CO_2_.

### Purification and Intoxication of RRSP-DT_B_ in CRC Cell Lines

Recombinant RRSP-DT_B_ and RRSP*-DT_B_ were expressed in *E. coli* BL21 (DE3) and purified over a HisTrap FF nickel affinity column as previously described (15). Eluted fractions were loaded onto a gravity column containing Strep-Tactin Superflow high capacity resin, followed by SUMO-tag removal and size exclusion purification over a Superdex 75 column using ÄKTA protein purifier purification system as previously described (15). For intoxication, CRC cell lines were seeded in 6-well plates (~70% confluency) overnight, after which medium was replaced with fresh medium containing either RRSP-DT_B_ or RRSP*-DT_B_ and incubated at indicated timepoints at 37°C in the presence of 5% CO_2_.

### Time-Lapse Video Microscopy

For RAS-less MEFs (6 x 10^3^ cells per well) were cultured in 96-well clear bottom white plates in corresponding complete growth medium and treated after 4 hours with RRSP-DT_B_ or RRSP*-DT_B_. Colorectal cancer cell lines were plated at ~80% confluency and cultured in 96-well clear bottom white plates. Complete growth medium with RRSP-DT_B_ or RRSP*-DT_B_ was added after overnight cell attachment. All cells were cultured were in Nikon Biostation CT and images were taken at indicated timepoints. Cell confluency was quantified using Nikon Elements software. IC50 concentrations were calculated using log(inhibitor) vs. response variable slope (four parameters) function in Graphpad Prism 8.

### Cell Viability and Cell Survival Assays

CRC cell lines were seeded in 96-well clear bottom white plates at ~80% confluency. Complete growth medium with RRSP-DT_B_ or RRSP*-DT_B_ was added after overnight cell attachment. After 72 hours, CellTiter-Glo (Promega) reagent was added to each well and luminescence was detected using Tecan Safire2 plate reader. For crystal violet assays, cells were treated as described above and were incubated for 48 hours. Following incubation cells were harvested and reseeded at low seeding densities in 6-well plates. Colony formation was monitored over 14 days, during which media was replaced every three days. On day 14 colonies were fixed in crystal violet fixing/staining solution (0.05% (g/vol) crystal violet, 1% formaldehyde, 1% (v/v) methanol in PBS. Open source ColonyArea ImageJ plug-in was used for quantitative analysis of the area % covered by the stained colonies (44). Due to high background from crystal violet staining in SW620 cells, stained wells were dissolved in 10% acetic acid and destained on rocker for 30 minutes. Absorbance was measured at 590 nm using Tecan Safire2 plate reader.]

### Proteome Human Phospho-Kinase Array

CRC cell lines were treated as described and washed in 1X PBS. Cells were solubilized using lysis buffer provided by the vendor (R&D Systems) and rocked for 30 minutes at 4°C. Suspension was spun for 5 minutes at 14,000 x *g* and supernatant was collected. Concentration of protein in the collected supernatant fluid determined using the BCA assay (ThermoFisher Scientific, no. 23227). 200 μg of sample lysate was applied to nitrocellulose membranes kinase arrays and incubated overnight at 4°C. Provided detection antibodies were incubated with specified concentrations as suggested by the supplier. Membrane arrays were acquired using Odyssey Infrared Imaging System (LI-COR Biosciences) and quantified by densitometry using NIH ImageJ software. Values from densitometry analysis were normalized to HSP60 control. Normalized value was then converted to Log2 fold change and plotted on heatmap using Graphpad Prism 8.

### Cell cycle flow cytometry

CRC cell lines were treated as described above. After 24 hours of treatment, cells were collected from medium, washed with 1X PBS, and released from well with Trypsin-EDTA (0.25%), phenol red (Invitrogen). Harvested cells were centrifuged at 700 x *g* for 5 minutes. Cells were washed twice in PBS and spun down at 700 x *g* for 5 minutes. PBS was removed and cells were resuspended in 600 μL of ice-cold PBS. Cell were permeabilized with addition of 1.4 mL of icecold ethanol slowly and incubated overnight at −20°C. Following two washes with PBS (centrifuged at 700 x *g* for 5 minutes), cells were stained in 200 μL PI staining solution (0.1% Triton X-100, 50 μg propidium iodide (BioLegend), 0.2 mg/mL RNase) for 30 minutes. Samples were analyzed on BD LSR Fortessa 1 Analyzer. At least 10,000 events were collected for each sample. Single cell populations were viewed and gated on cyanine-3 area (Cy3-A) versus cyanine-3 width (Cy3-W) channels, to eliminate doublet events. ModFit LT Software (Version 5) was used for cell cycle analysis.

### Disclosure of Potential Conflicts of Interest

K.J.F.S. has been granted a patent (US 10,829,752 B2) on use of RRSP to treat cancer. K.J.F.S. is a consultant for Buoy Health on topics unrelated to this manuscript. K.J.F.S. has a significant financial interest in Situ Biosciences, LLC, a contract research organization that pursues research unrelated to cancer. C.K.S. is an intern as a scientific and financial advisor for Aspire Capital Partners, LLC, which invests in oncotherapies.

## Authors’ Contributions

C.K.S. designed and conducted all experiments and wrote the manuscript. M.B. conducted preliminary experiments with the Ras-less MEF cells. V.V. contributed reagents, assisted with experimental design, and edited the manuscript. K.J.F.S. oversaw all aspects of design, conduct, and analysis of the experiments and edited the manuscript.

## Acknowledgments

We thank the Frederick National Laboratory for Cancer Research (FNLCR) for providing the RAS-less MEF cells, KRAS mutant cell lines, and hybridoma cells for the pan-RAS antibody directed against the G-domain. Matthew Kieffer and all members of the Satchell laboratory are thanked for valuable intellectual input and technical support. This work was funded by grants from the Chicago Biomedical Consortium, H Foundation, Northwestern Medicine Catalyst Fund, and National Institute of Health R01 AI092825 (to K.J.F.S). C.K.S. was supported by a fellowship from the National Cancer Institute (T32 CA09560). High content imaging was performed on the Nikon Biostation CT system purchased with the support of NIH grant S10 OD021704. Flow cytometry core services were provided by the Northwestern University RHLCCC Flow Cytometry Facility supported by NIH grant P30 CA060553.

## Footnotes

**Note:** Supplementary data for this article (Suppl. Fig. 1–6)

---

**Supplementary Figure 1.**
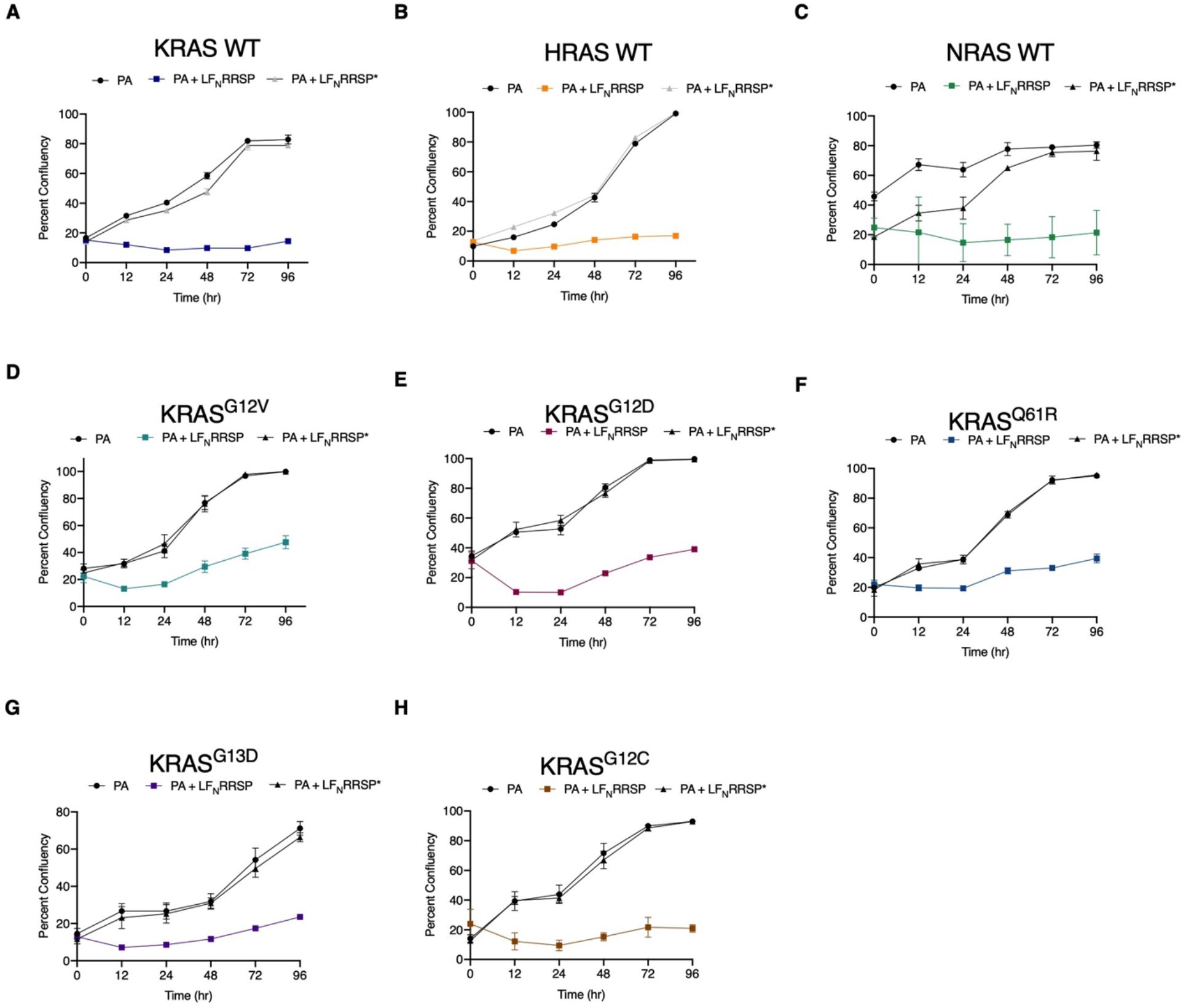
Growth inhibition of RAS-less MEFs cell lines. (**A-H**) Growth inhibition observed in of RAS-less MEF cell lines at indicted timepoints following treatment with PA alone or in combination with LF_N_RRSP or LF_N_RRSP*; *n* = 3. Results are expressed as ± SEM.

**Supplementary Figure 2.**
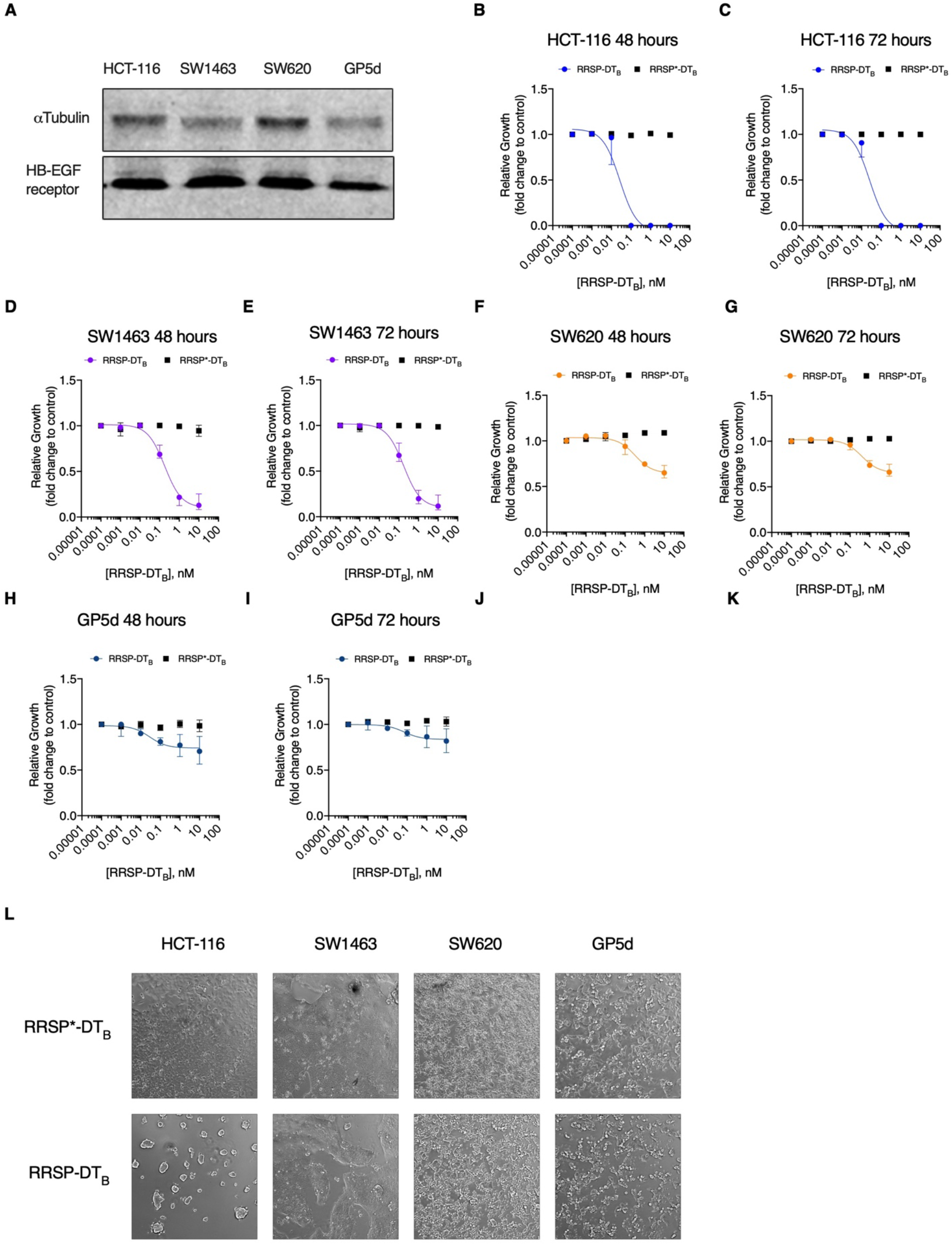
Growth inhibition in RRSP-DT_B_ treated CRC cell lines. (**A**). Western blot analysis of HB-EGF receptor (DT receptor) from untreated CRC cell line lysates. Vinculin was used as loading control. (**B-K**) Fitted dose response curve of RRSP-DT_B_ in CRC cell lines after 48 and 72 hours. Results are displayed as mean ± SEM, *n* = 4. (**L**) Representative brightfield images of CRC cell lines treated with either RRSP-DT_B_ or RRSP*-DT_B_ (0.1 nM) after 24 hours.

**Supplementary Figure 3.**
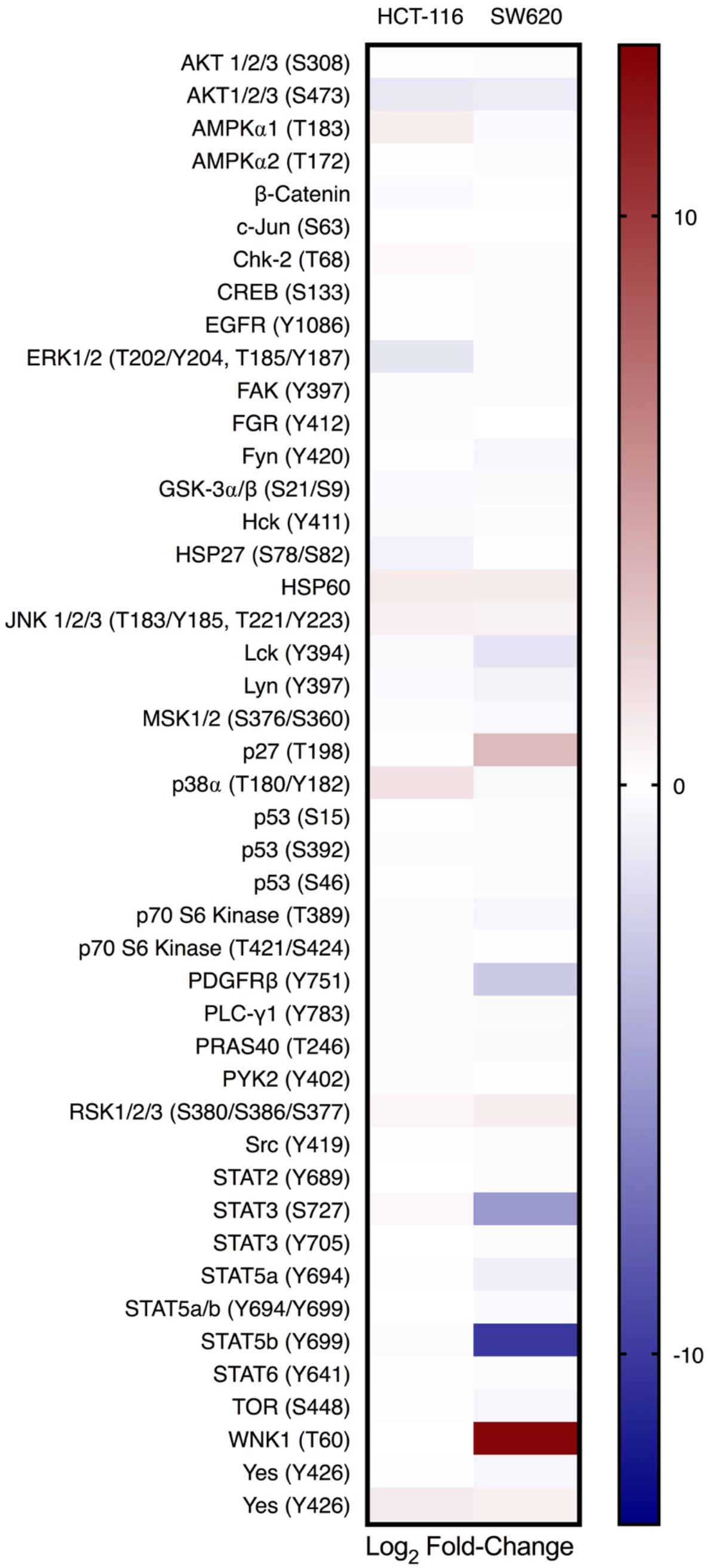
Heatmap of human phosphor-kinase array in HCT-116 and SW620 cells treated RRSP-DT_B_ treated samples. Densitometric analysis of phospho-kinase array depicted through a heatmap of relative phosphorylated proteins levels in response to 10 nM RRSP-DT_B_ compared to PBS control in HCT-116 and SW620 cells after 24 hr.

**Supplementary Figure 4.**
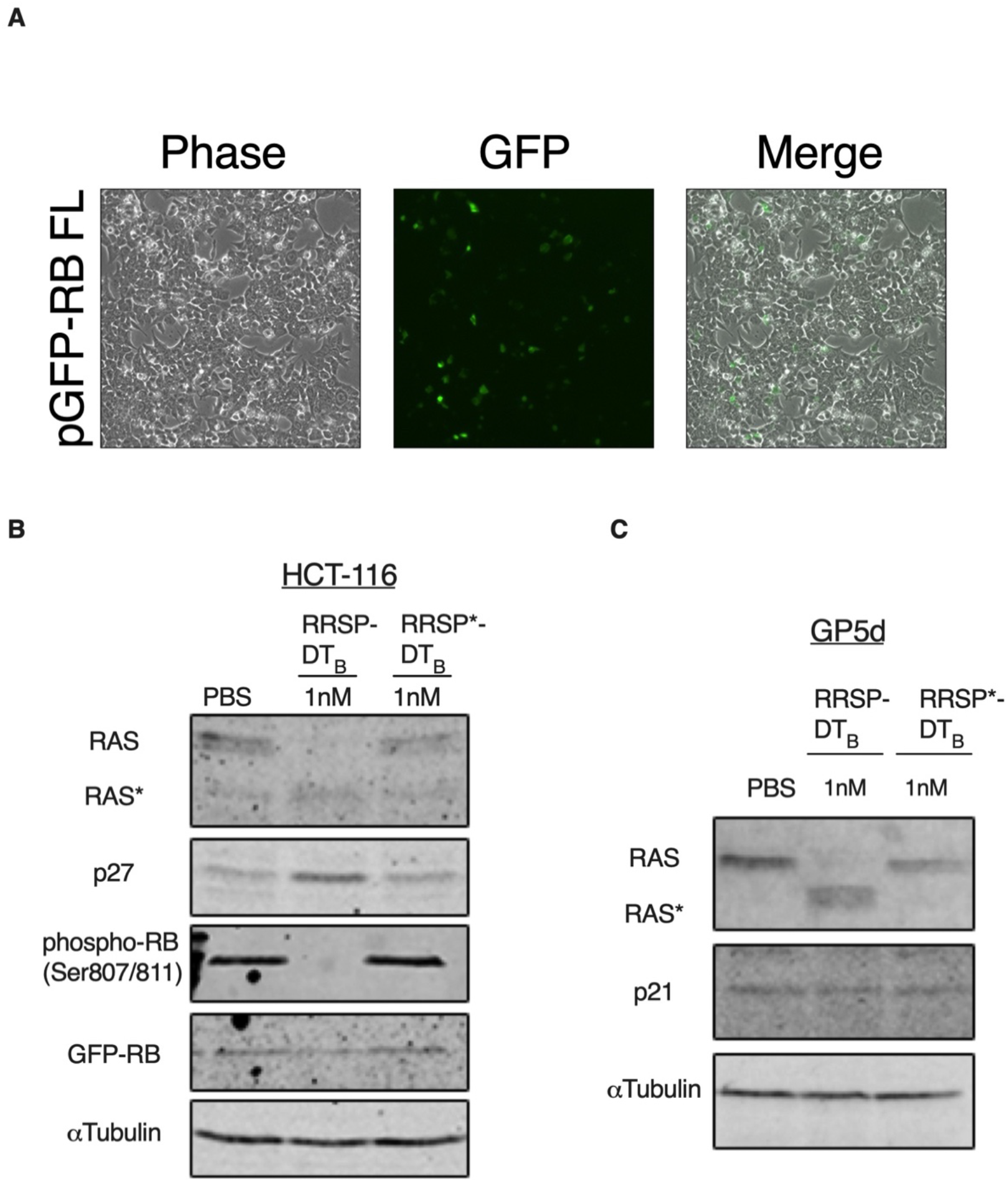
Total RB expression in CRC cell lines. (**A**) Representative images of HCT-116 cells transfected with pGFP-RB after 24 hr (**B**) Western blot analysis of HCT-116 cells transfected with GFP tagged RB following treatment with either PBS, RRSP-DT_B_ or RRSP*-DT_B_ after 24 hours. Total RB was detected using anti-GFP primary antibody. aTubulin was used as a gel loading control. (**C**) Western blot analysis of p21 levels in GP5d cell treated with either PBS, RRSP-DT_B_ or RRSP*-DT_B_ after 24 hr.

**Supplementary Figure 5.**
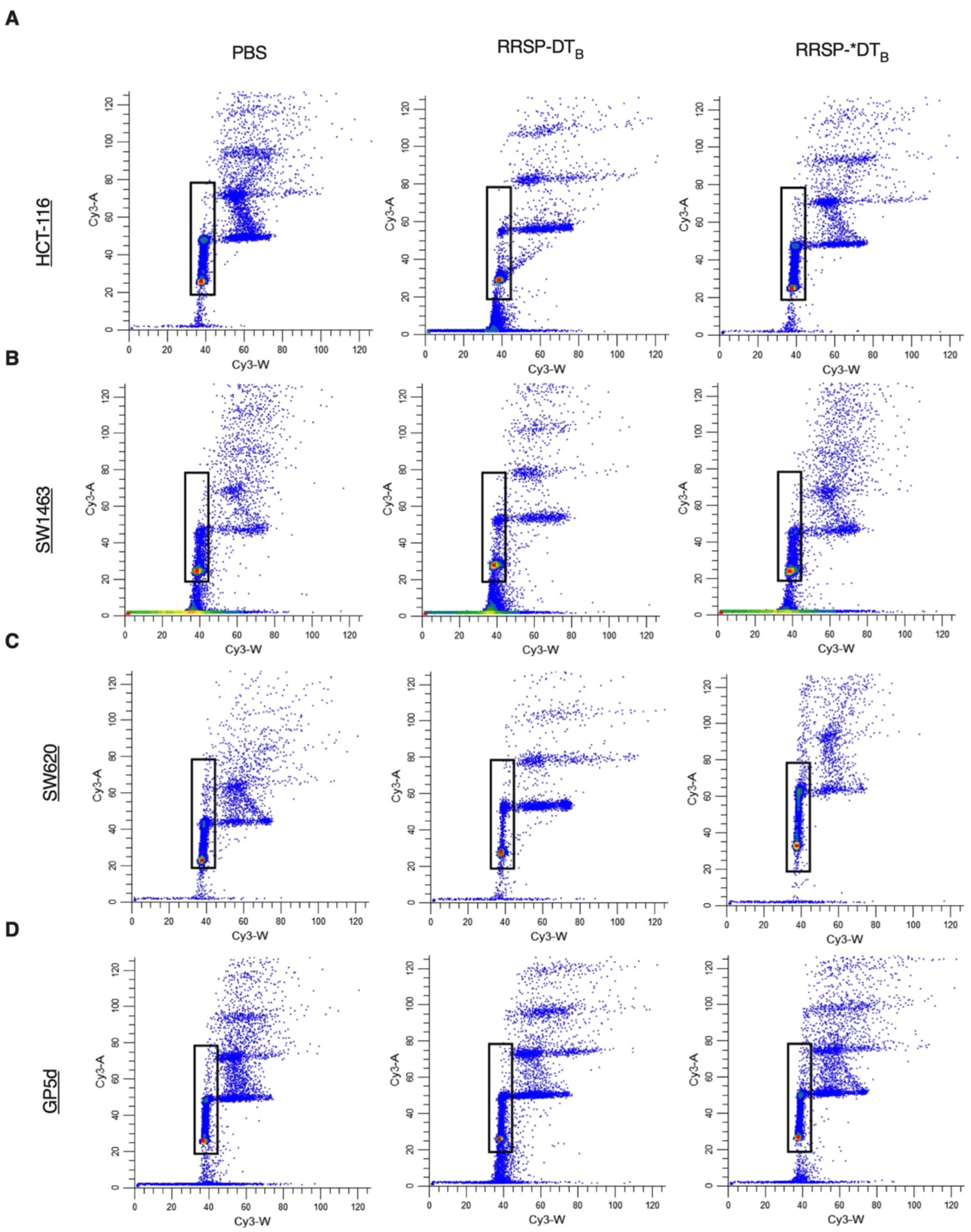
Cell cycle analysis of CRC cell lines treated RRSP-DT_B_. (**A-D**) Representative flow cytometry plots of CRC cell lines treated with either PBS, RRSP-DT_B_ or RRSP*-DT_B_ (1 nM) after 24 hours. Gating parameters were used to only collect single live cell populations.

**Supplementary Figure 6.**
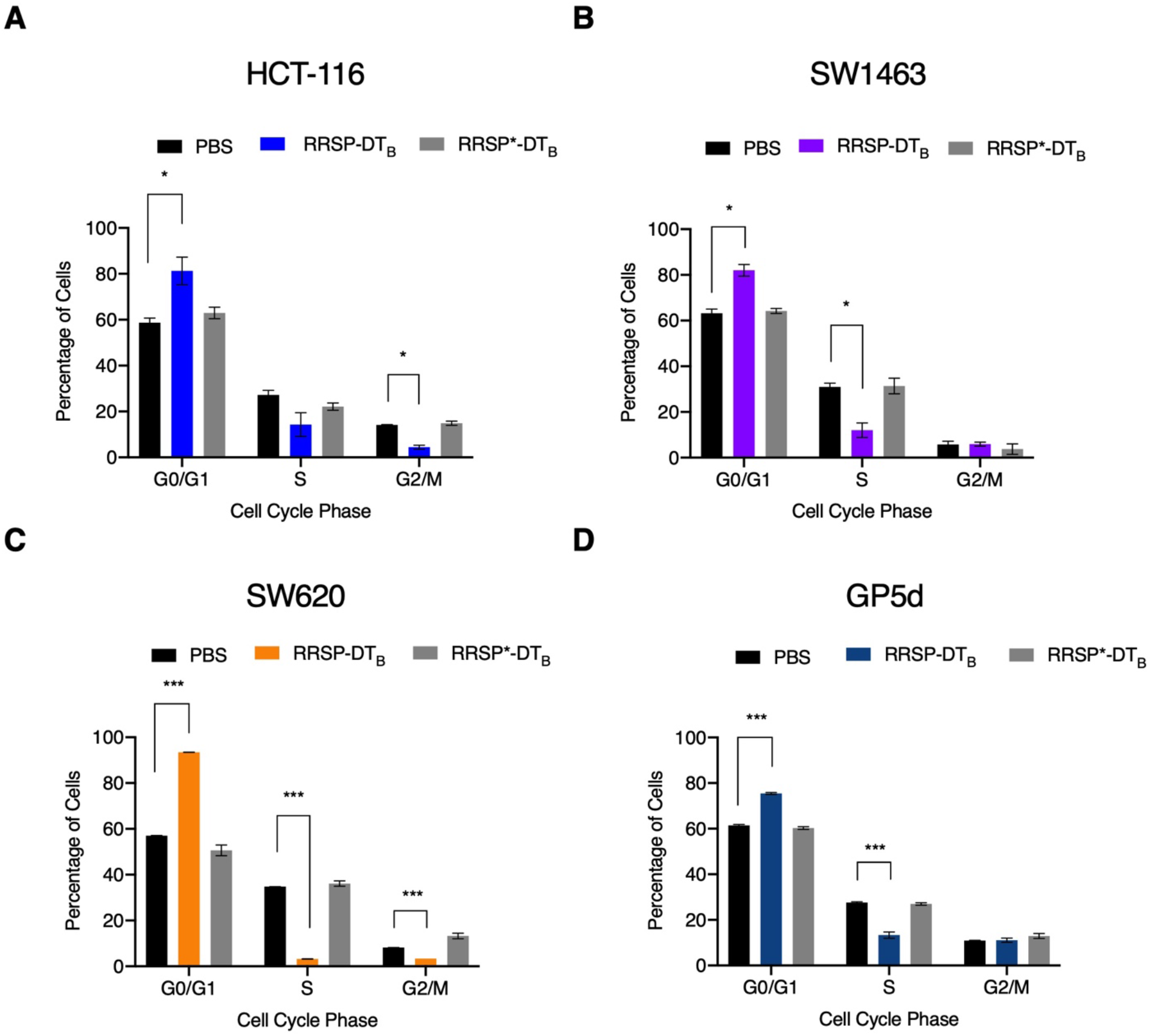
Cell cycle analysis of RRSP-DT_B_ treated CRC cell lines. (**A-D**) Quantitative analysis of cell cycle analysis of CRC cell lines treated with PBS, 1 nM RRSP-DT_B_ or 1 nM RRSP*-DT_B_ for 24 hours; *n* = 3. Results are expressed as means ± SEM of three independent experiments (*P < 0.05, **P<0.01, ****P<0.0001 versus PA control as determined through one-way ANOVA followed by Dunnett’s multiple comparison test).

